# BO-LSTM: Classifying relations via long short-term memory networks along biomedical ontologies

**DOI:** 10.1101/336719

**Authors:** Andre Lamurias, Luka A. Clarke, Francisco M. Couto

## Abstract

Recent studies have proposed deep learning techniques, namely recurrent neural networks, to improve biomedical text mining tasks. However, these techniques rarely take advantage of existing domain-specific resources, such as ontologies. In Life and Health Sciences there is a vast and valuable set of such resources publicly available, which are continuously being updated. Biomedical ontologies are nowadays a mainstream approach to formalize existing knowledge about entities, such as genes, chemicals, phenotypes, and disorders. These resources contain supplementary information that may not be yet encoded in training data, particularly in domains with limited labeled data.

We propose a new model, BO-LSTM, that takes advantage of domain-specific ontologies, by representing each entity as the sequence of its ancestors in the ontology. We implemented BO-LSTM as a recurrent neural network with long short-term memory units and using an open biomedical ontology, which in our case-study was Chemical Entities of Biological Interest (ChEBI). We assessed the performance of BO-LSTM on detecting and classifying drug-drug interactions in a publicly available corpus from an international challenge, composed of 792 drug descriptions and 233 scientific abstracts. By using the domain-specific ontology in addition to word embeddings and WordNet, BO-LSTM improved both the F1-score of the detection and classification of drug-drug interactions, particularly in a document set with a limited number of annotations. Our findings demonstrate that besides the high performance of current deep learning techniques, domain-specific ontologies can still be useful to mitigate the lack of labeled data.

**Author summary:** A high quantity of biomedical information is only available in documents such as scientific articles and patents. Due to the rate at which new documents are produced, we need automatic methods to extract useful information from them. Text mining is a subfield of information retrieval which aims at extracting relevant information from text. Scientific literature is a challenge to text mining because of the complexity and specificity of the topics approached. In recent years, deep learning has obtained promising results in various text mining tasks by exploring large datasets. On the other hand, ontologies provide a detailed and sound representation of a domain and have been developed to diverse biomedical domains. We propose a model that combines deep learning algorithms with biomedical ontologies to identify relations between concepts in text. We demonstrate the potential of this model to extract drug-drug interactions from abstracts and drug descriptions. This model can be applied to other biomedical domains using an annotated corpus of documents and an ontology related to that domain to train a new classifier.

## Introduction

Traditional relation extraction methods employ machine learning algorithms, often using kernel functions in conjunction with Support Vector Machines [1, 2] or based on features extracted from the text [3]. In recent years, deep learning techniques have obtained promising results in various Natural Language Processing (NLP) tasks [4], including relation extraction. These techniques have the advantage of being easily adaptable to multiple domains, using models pre-trained on unlabeled documents [5]. The success of deep learning for text mining is in part due to the high quantity of raw data available and the development of word vector models such as Word2vec [6] and GloVe [7]. These models can use unlabeled data to predict the most probable word according to the context words (or vice-versa), leading to meaningful vector representations of the words in a corpus, known as word embeddings.

A high volume of biomedical information relevant to the detection of Adverse Drug Reactions (ADRs), such as Drug-Drug Interactions (DDI), is mainly available in articles and patents [8] A recent review of studies about the causes of hospitalization in adult patients has found that ADRs were the most common cause, accounting for 7% of hospitalizations [9]. Another systematic review focused on the European population, identified that 3.5% of hospital admissions were due to ADRs, while 10.1% of the patients experienced ADRs during hospitalization [10].

The knowledge encoded in the ChEBI (Chemical Entities of Biological Interest) ontology is highly valuable to detect and classify DDIs, since we not only get access to important characteristics of each individual compound but more importantly we gain access to the underlying semantics of the relations between compounds. For instance, dopamine (CHEBI:18243), a chemical compound with several important roles in the brain and body, can be characterized as being a catecholamine (CHEBI:33567), an aralkylamino compound (CHEBI:64365) and an organic aromatic compound (CHEBI:33659) (Fig 1). When predicting if a certain drug interacts with dopamine, its ancestors will provide additional information that is not usually directly expressed in the text. While the reader can consult additional materials to better understand a biomedical document, traditional relation extraction models are trained solely on features extracted from the training corpus. Thus, ontologies are an advantage to relation extraction models due to the semantics encoded in them regarding a particular domain. Since ontologies are described in a common machine-readable format, methods based on ontologies can be applied to different domains and incorporated with other sources of knowledge, bridging the semantic gap between relation extraction models, data sources, and results [11].

**Fig 1.**
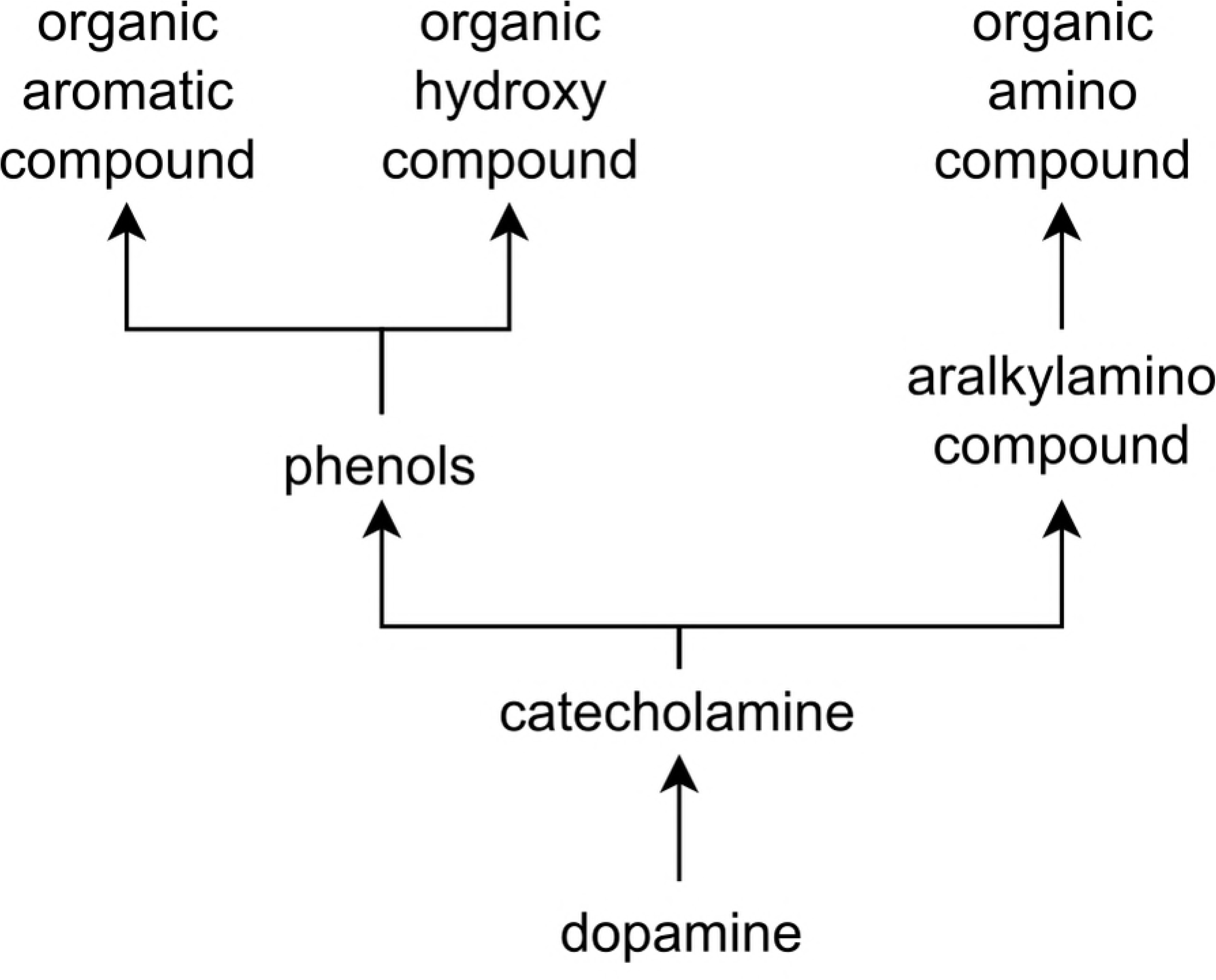
An excerpt of the ChEBI ontology showing the first ancestors of dopamine, using “is-a” relationships.

### Deep learning for biomedical NLP

Current state-of-the-art text mining methods employ deep learning techniques, such as Recurrent Neural Networks (RNN), to train classification models based on word embeddings and other features. These methods use architectures composed of multiple layers, where each layer attempts to learn a different kind of representation of the input data. This way, different types of tasks can be trained using the same input data. Furthermore, there is no need to manually craft features for a specific task.

Long Short-Term Memory (LSTM) networks have been proposed as an alternative to regular RNN [12]. LSTMs are a type of RNN that can handle long dependencies, and thus are suitable for NLP tasks, which involve long sequences of words. When training the weights of an RNN, the contribution of the gradients may vanish while propagating for long sequences of words. LSTM units account for this vanishing gradient problem through a gated architecture, which makes it easier for the model to capture long-term dependencies. Recently, LSTMs have been applied to relation extraction tasks in various domains. Miwa and Bansal [13] presented a model that extracted entities and relations based on bidirectional tree-structured and sequential LSTM-RNNs. The authors evaluated this model on three datasets, including the SemEval 2010 Task 8 dataset, which defines 10 general semantic relations types between nominals.

Bidirectional LSTMs have been proposed for relation extraction, obtaining better results than one-directional LSTMs on the SemEval 2010 dataset [14]. In this case, at each time step, there are two LSTM layers, one that reads the sentence from left to right, and another that reads from right to left. The output of both layers is combined to produce a final score.

The model proposed by Xu et al. [15] combines Shortest Dependency Paths (SDP) between two entities in a sentence with linguistic information. SDPs are informative features for relations extraction since these contain the words of the sentence that refer directly to both entities. This model has a multichannel architecture, where each channel makes use of information from a different source along the SDP. The main channel, which contributes the most to the performance of the model, uses word embeddings trained on the English Wikipedia with Word2vec. Additionally, the authors study the effect of adding channels consisting of the part-of-speech tags of each word, the grammatical relations between the words of the SDP, and the WordNet hypernyms of each word. Using all four channels, the F1-score of the SemEval 2010 Task 8 was 0.0135 higher than when using only the word embeddings channel. Although WordNet can be considered an ontology, its semantic properties were not explored in this work, since only the class of word is extracted, and the relations between classes are not explored.

### Ontologies for biomedical text mining

While machine learning classifiers trained on word embeddings can learn to detect relations between entities, these classifiers may miss the underlying semantics of the entities according to their respective domain. However, the semantics of a given domain are, in some cases, available in the form of an ontology. Ontologies aim at providing a structured representation of the semantics of the concepts in a domain and their relations. In this paper, we consider a domain-specific ontology as a directed acyclic graph where each node is a concept (or entity) of the domain and the edges represent known relations between these concepts [16]. This is the traditional representation of existing biomedical ontologies, which are nowadays a mainstream approach to formalize knowledge about entities, such as genes, chemicals, phenotypes, and disorders.

Biomedical ontologies are usually publicly available and cover a large variety of topics related to Life and Health Sciences. In this paper, we use ChEBI, an ontology for chemical compounds with biological interest, where each node corresponds to a chemical compound [17]. The latest release of ChEBI contains nearly 54k compounds and 163k relationships. Note that, the success of exploring a given biomedical ontology for performing a specific task can be easily extended to other topics due to the common structure of biomedical ontologies. For example, the same measures of metadata quality have been successfully applied to resources annotated with different biomedical ontologies [18].

Other authors have previously combined ontological information with neural networks, to improve the learning capabilities of a model. Li et al. [19] mapped each word to a WordNet sense disambiguation to account for the different meanings that a word may have and the relations between word senses. Ma et al. [20] proposed the LSTM-OLSI model, which indexes documents based on the word-level contextual information from the DBpedia ontology and document-level topic modeling. Some authors have explored graph embedding techniques, converting relations to a low dimensional space which represents the structure and properties of the graph [21]. For example, Kong et al. [22] combined heterogeneous sources of information, such as ontologies, to perform multi-label classification, while Dasigi et al. [23] presented an embedding model based on ontology concepts to represent word tokens.

However, few authors have explored biomedical ontologies for relation extraction. Textpresso is a project that aims at helping database curation by automatically extracting biomedical relations from research articles [24]. Their approach incorporates an internal ontology to identify which terms may participate in relations according to their semantics. Other approaches measure the similarity between the entities and use the value as a feature for a machine learning classifier [25]. One of the teams that participated in the BioCreative VI ChemProt task used ChEBI and Protein Ontology to extract additional features for a neural network model that extracted relation between chemicals and proteins [26]. To the best of our knowledge, our work is the first attempt at incorporating ancestry information from biomedical ontologies with deep learning to extract relations from text.

In this manuscript, we propose a new model, BO-LSTM that can explore domain information from ontologies to improve the task of biomedical relation extraction using deep learning techniques. The code and results obtained with the model can be found on our GitHub repository (https://github.com/lasigeBioTM/BOLSTM), while a Docker image is also available (https://hub.docker.com/r/andrelamurias/bolstm), simplifying the process of training new classifiers and applying them to new data. We compare the effect of using ChEBI, a domain-specific ontology, and Wordnet, a generic English language ontology, as external sources of information to train a classification model based on LSTM networks. This model was evaluated on a publicly available corpus of 792 drug descriptions and 233 scientific abstracts annotated with DDIs relevant to the study of adverse drug effects. Using the domain-specific ontology in addition to word embeddings and WordNet, BO-LSTM improved the F1-score of the classification of DDIs by 0.0207. Our model was particularly efficient with document types that were less represented in the training data. These results validate our hypothesis that domain-specific information is useful to complement data-intensive approaches such as deep learning.

## Results

We evaluated the performance of our BO-LSTM model on the SemEval 2013: Task 9 DDI extraction corpus [27]. This gold standard corpus consists of 792 texts from DrugBank [], describing chemical compounds, and 233 abstracts from the MedLine database [28]. DrugBank is a cheminformatics database containing detailed drug and drug target information, while MedLine is a database of bibliographic information of scientific articles in Life and Health Sciences. Each document was annotated with pharmacological substances and sentence-level DDIs. We refer to each combination of entities mentioned in the same sentence as a candidate pair, which could either be positive if the text describes a DDI, or negative otherwise. In other words, a negative candidate is a candidate pair that is not described as interacting in the text. Each positive DDI was assigned one of four possible classes: mechanism, effect, advice, and int, when none of the others were applicable.

In the context of the competition, the corpus was separated into training and testing sets, containing both DrugBank and MedLine documents. After shuffling we used 80% of the training set to train the model and 20% as a validation set. This way, the validation set contained both DrugBank and MedLine documents, and overfitting to a specific document type is avoided. It has been shown that the DDIs of the MedLine documents are more difficult to detect and classify, with the best systems having almost a 30 point F1-score difference to the DrugBank documents [29].

We implemented the BO-LSTM model in Keras, a Python-based deep learning library, using the Tensorflow backend. The overall architecture of the BO-LSTM model is presented in Fig 2. More details about each layer can be found in the Methods section. We focused on the effect of using different sources of information to train the model. As such, we tuned the hyperparameters to obtain reasonable results, using as reference the values provided by other authors that have applied LSTMs to this gold standard [30, 31]. We first trained the model using only the word embeddings of the SDP of each candidate pair (Fig 2A). Then we tested the effect of adding the Wordnet classes as a separate embedding and LSTM layer (Fig 2B) Finally, we tested two variations of the ChEBI channel: first using the concatenation of the sequence of ancestors of each entity (Fig 2C), and second using the sequence of common ancestors of both entities (Fig 2D).

**Fig 2.**
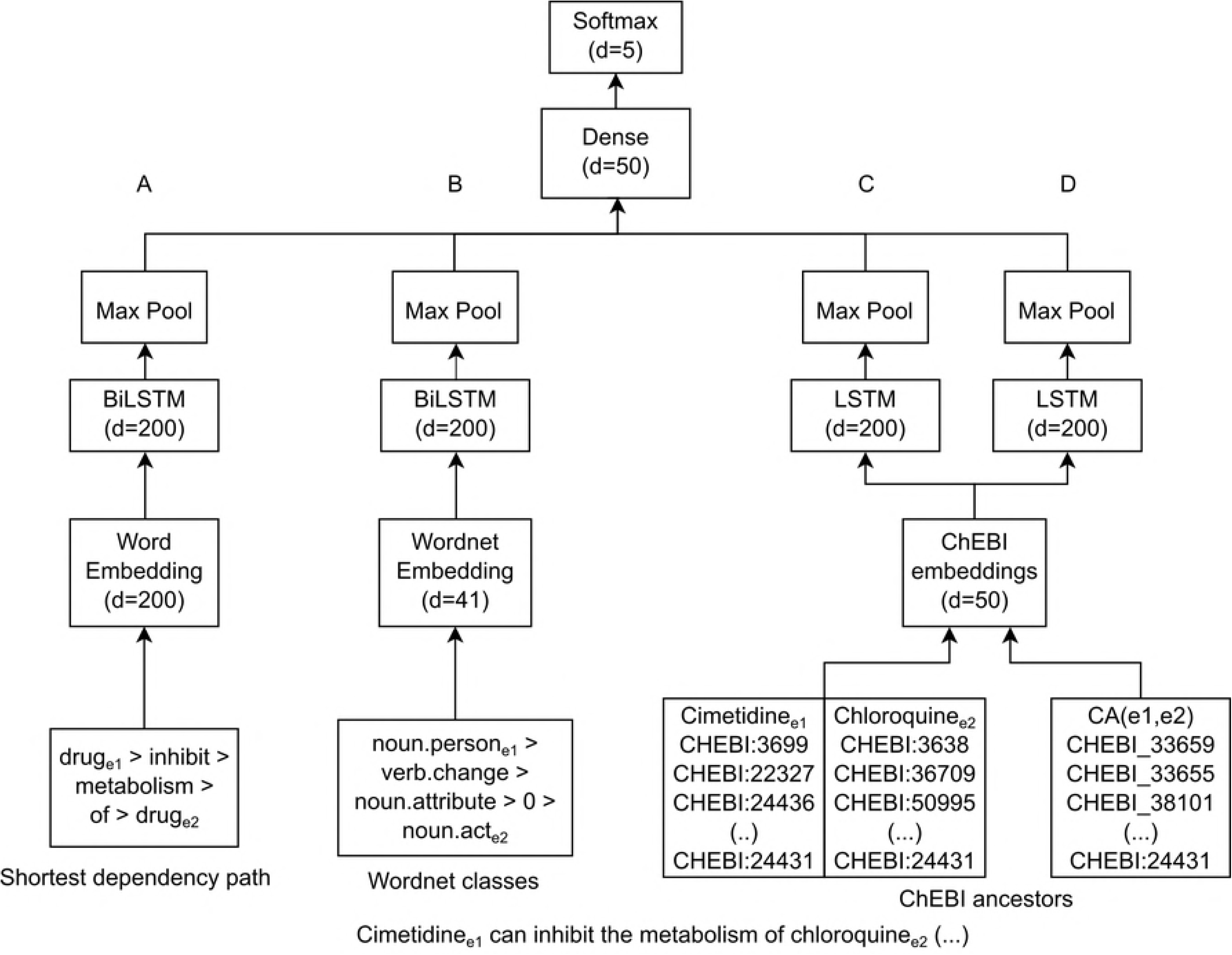
BO-LSTM Model architecture, using a sentence from the Drug-Drug Interactions corpus as an example. Each box represents a layer, with an output dimension, and merging lines represent concatenation. We refer to (A) as the Word embeddings channel, (B) the Wordnet channel and (C) the ancestors concatenation channel and (D) the common ancestors channel.

Table 1 shows the DDI detection results obtained with each configuration using the evaluation tool provided by the SemEval 2013: Task 9 organizers on the gold standard, while Table 2 shows the DDI classification results, using the same evaluation tool and gold standard. The difference between these two tasks is that while detection ignores the type of interactions, the classification task requires identifying the positive pairs and also their correct interaction type. We compare the performance on the whole gold standard, and on each document type (DrugBank and MedLine). The first row of each table shows the results obtained using an LSTM network trained solely on the word embeddings of the SDP of each candidate pair. Then, we studied the impact of adding each information channel on the performance of the model, and the effect of using all information channels, as shown in Fig 2.

**Table 1.**
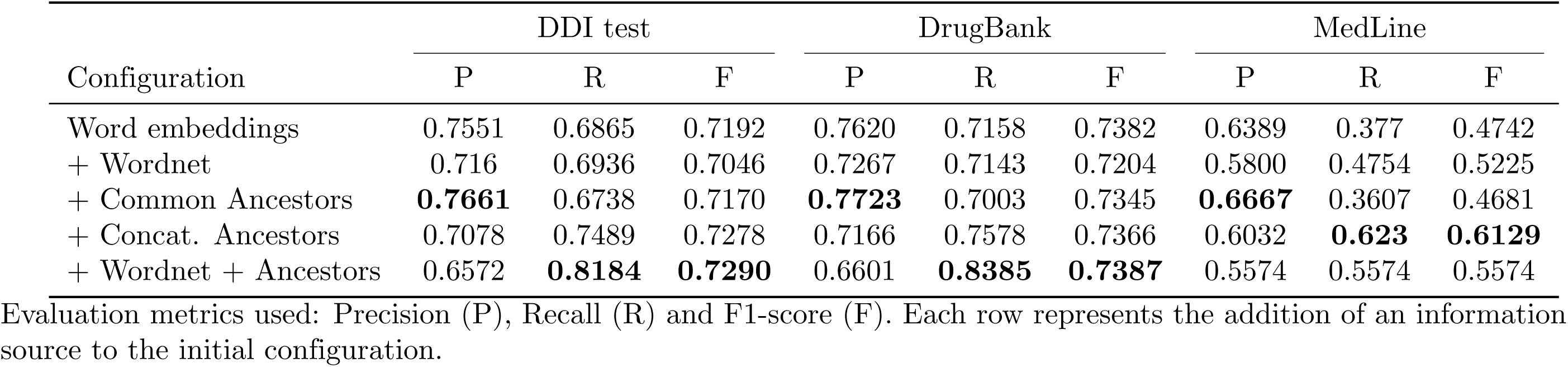
Evaluation scores obtained for the DDI detection task on the DDI corpus and on each type of document, comparing different configurations of the model.

| Configuration | DDI test |  |  | DrugBank |  |  | MedLine |  |  |
| --- | --- | --- | --- | --- | --- | --- | --- | --- | --- |
|  | P | R | F | P | R | F | P | R | F |
| Word embeddings | 0.7551 | 0.6865 | 0.7192 | 0.7620 | 0.7158 | 0.7382 | 0.6389 | 0.377 | 0.4742 |
| + Wordnet | 0.716 | 0.6936 | 0.7046 | 0.7267 | 0.7143 | 0.7204 | 0.5800 | 0.4754 | 0.5225 |
| + Common Ancestors | **0.7661** | 0.6738 | 0.7170 | **0.7723** | 0.7003 | 0.7345 | **0.6667** | 0.3607 | 0.4681 |
| + Concat. Ancestors | 0.7078 | 0.7489 | 0.7278 | 0.7166 | 0.7578 | 0.7366 | 0.6032 | **0.623** | **0.6129** |
| + Wordnet + Ancestors | 0.6572 | **0.8184** | **0.7290** | 0.6601 | **0.8385** | **0.7387** | 0.5574 | 0.5574 | 0.5574 |

Evaluation metrics used: Precision (P), Recall (R) and F1-score (F). Each row represents the addition of an information source to the initial configuration.

**Table 2.** Evaluation scores obtained for the DDI classification task on the DDI corpus and on each type of document, comparing different configurations of the model.

| Configuration | DDI test |  |  | DrugBank |  |  | MedLine |  |  |
| --- | --- | --- | --- | --- | --- | --- | --- | --- | --- |
|  | P | R | F | P | R | F | P | R | F |
| Word embeddings | 0.5819 | 0.5291 | 0.5542 | 0.5868 | 0.5512 | 0.5685 | 0.5000 | 0.2951 | 0.3711 |
| + Wordnet | 0.5754 | 0.5574 | 0.5663 | 0.5845 | 0.5745 | **0.5795** | 0.4600 | 0.3770 | 0.4144 |
| + Common Anc. | **0.5968** | 0.5248 | 0.5585 | **0.6045** | 0.5481 | 0.5749 | **0.5152** | 0.2787 | 0.3617 |
| + Concat. Anc. | 0.5282 | 0.5589 | 0.5431 | 0.5286 | 0.5590 | 0.5434 | 0.4921 | **0.5082** | **0.5000** |
| + Wordnet + Anc. | 0.5182 | **0.6454** | **0.5749** | 0.5171 | **0.6568** | 0.5787 | 0.4590 | 0.4590 | 0.4590 |

Evaluation metrics used: Precision (P), Recall (R) and F1-score (F). Each row represents the addition of an information source to the initial configuration.

For the detection task, using the concatenation of ancestors results in an improvement of the F1-score in the MedLine dataset, contributing to an overall improvement of the F1-score in the full test set. The most notable improvement was in the recall of the MedLine dataset, where the concatenation of ancestors increased this score by 0.246. The usage of ontology ancestors did not improve the F1-score of detection of DDIs in the DrugBank dataset. In every test set, it is possible to observe that the concatenation of ancestors results in a higher recall, while considering only the common ancestors is more beneficial to precision. Combining both approaches with the Wordnet channel results in a higher F1-score.

Regarding the classification task (Table 2), the F1-score was improved on each dataset by the usage of the ontology channel. Considering only the common ancestors led to an improvement of the F1-score in the DrugBank dataset and on the full corpus, while the concatenation improved the MedLine F1-score, similarly to the detection results.

To better understand the contribution of each channel, we studied the relations detected by each configuration by one or more channels, and which of those were also present in the gold standard. Fig 3 and Fig 4 show the intersection of the results of each channel in the full, DrugBank, and MedLine test sets. We compare only the results of the detection task, as it is simpler to analyze and show the differences in results of different configurations. In Fig 3, we can visualize false negatives as the number of relations unique to the gold standard and the false positives of each configuration as the number of relations that does not intersect with the gold standard. The difference between the values of this figure and the sum of their respective values in Fig 4 is due to the system being executed once for each dataset. Overall 369 relations in the full test set were not detected by any configuration of our system, out of a total of 979 relations in the gold standard. We can observe that 60 relations were detected only when adding the ontology channels.

**Fig 3.**
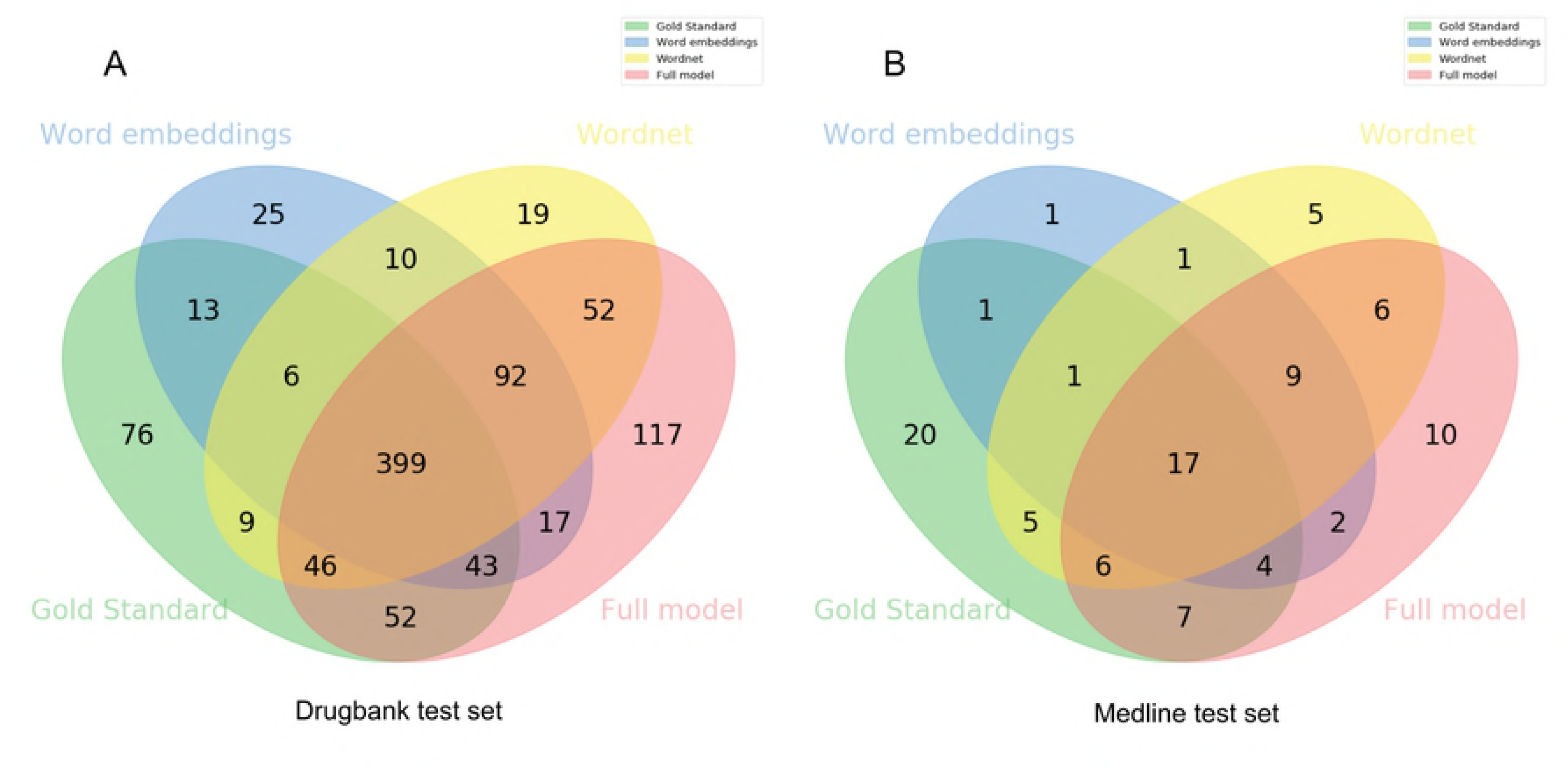
Venn diagram demonstrating the contribution of each configuration of the model to the results of the full test set. The intersection of each channel with the gold standard represents the number of true positives of that channel, while the remaining correspond to false negatives and false positives.

**Fig 4.**
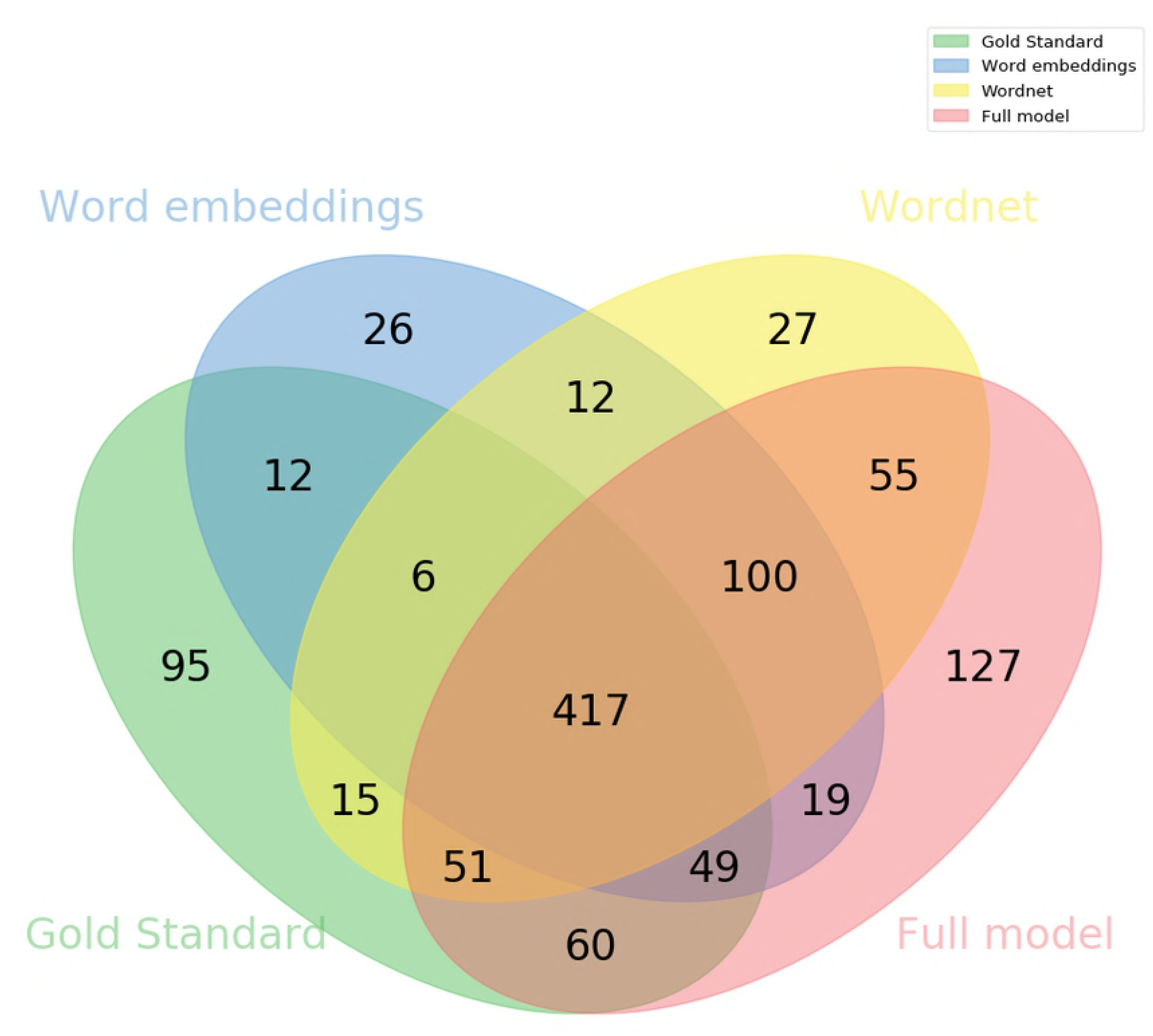
Venn diagram demonstrating the contribution of each configuration of the model to the DrugBank (A) and MedLine (B) test set results. The intersection of each channel with the gold standard represents the number of true positives of that channel, while the remaining correspond to false negatives and false positives.

In the MedLine test set, the ontology channel identified 7 relations that were not identified by any other configuration (Fig 4B). One of these relations was the effect of quinpirole treatment on amphetamine sensitization. Quinpirole has 27 ancestors in the ChEBI ontology, while amphetamine has 17, and they share 10 of these ancestors, with the most informative being “organonitrogen compound”. While this information is not described in the original text, but only encoded in the ontology, it is relevant to understand if the two entities can participate in a relation. However, this comes at the cost of precision, since 10 incorrect DDIs were classified by this configuration.

## Discussion

Comparing the results across the two types of documents, we can observe that our model was most beneficial to the MedLine test set. This set contains only 1301 sentences from 142 documents for training, while the DrugBank set contains 5675 sentences from 572 documents. Naturally, the patterns of the DrugBank documents will be easier to learn than the ones of the MedLine documents because more examples are shown to the model. Furthermore, the MedLine set has 0.18 relations per sentence, while the DrugBank set has 0.67 relations per sentence. This means that DDIs are described much more sparsely than in the DrugBank set. This demonstrates that our model is able to obtain useful knowledge that is not described in the text.

One disadvantage of incorporating domain information in a machine learning approach is that it reduces its applicability to other domains. However, biomedical ontologies have become ubiquitous in biomedical research. One of the most successful cases of a biomedical ontology is the Gene Ontology, maintained by the Gene Ontology Consortium [32]. The Gene Ontology defines over 40,000 concepts used to describe the properties of genes. This project is constantly updated, with new concepts and relations being added every day. However, there are ontologies for more specific subjects, such as microRNAs [33], radiology terms [34] and rare diseases [35]. BioPortal is a repository of biomedical ontology, currently hosting 685 ontologies. Furthermore, while manually labeled corpora are created specifically to train and evaluate text mining applications, ontologies have diverse applications, i.e., they are not developed for this specific purpose.

We evaluate the proposed model on the DDI corpus because it is associated with a SemEval task, and for this reason, it has been the subject of many studies since its release. However, while applying our model to a single domain, we designed its architecture so it can fit any other domain-specific ontology. In fact, the methodology proposed can be easily followed to apply to any other biomedical ontology that describes the concepts of a particular domain. For example, the Disease Ontology [36], that describes relations between human diseases, could be used with the BO-LSTM model on a disease relation extraction task, as long as there is an annotated training corpus.

While we studied the potential of domain-specific ontologies based only on the ancestors of each entity, there are other ways to integrate semantic information from ontologies into neural networks. For example, one could consider only the ancestors with highest information content, since those would be the most helpful to characterize an entity. The information content can be estimated either by the probability of a given term in the ontology or in an external dataset. Alternatively, a semantic similarity measure that accounts for non-transitive relations could be used to find similar concepts to the entities of the relation [37], or one that considers only the most relevant ancestors [38].

## Methods

In this section, we describe the proposed BO-lSTM model in detail, as shown in Fig 2, with a focus on the aspects that refer to the use of biomedical ontologies.

### Data preparation

The objective of our work is to identify and classify relations between biomedical entities found in natural language text. We assume that the relevant entities are already recognized. Therefore, we process the input data in order to generate instances to be classified by the model. Considering the set of entities *E* mentioned in a sentence, we generate 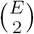 instances of that sentence. We refer to each instance as a candidate pair, identified by the two entities that constitute that pair, regardless of the order. A relation extraction model will assign a class to each candidate pair. In some cases, it is enough to simply classify the candidate pairs as negative or positive, while in other cases different types of positive relations are considered.

An instance should contain the information necessary to classify a candidate pair. Therefore, after tokenizing each sentence, we obtain the Shortest Dependency Path (SDP) between the entities of the pair. For example, in the sentence “Laboratory Tests Response to Plenaxis_e1_ should be monitored by measuring serum total testosterone_e1_ concentrations just prior to administration on Day 29 and every 8 weeks thereafter”, the shortest path between the entities would be Plenaxis - Response - monitored - by - measuring - concentrations - testosterone. For both tokenization and dependency parsing, we use the spacy software library (https://spacy.io/). The text of each entity that appears in the SDP, including the candidate entities, is replaced by the generic string to reduce the effect of specific entity names on the model. For each element of the SDP, we obtain the WordNet hypernym class using the tool developed by Ciaramita and Altun [39].

To focus our attention on the effect of the ontology information, we use pre-trained word embedding vectors. Pyysalo et al. [40] released a set of vectors trained on PubMed abstracts (nearly 23 million) and PubMed Central full documents (nearly 700k), trained with the word2vec algorithm [6]. Since these vectors were trained on a large biomedical corpus, it is likely that its vocabulary will contain more words relevant to the biomedical domain than the vocabulary of a generic corpus.

We match each entity to an ontology concept so that we can then obtain its ancestors. Ontology concepts contain an ID, a preferred label, and, in most cases, synonyms. While preprocessing the data, we match each entity to the ontology using fuzzy matching. The adopted implementation uses the Levenshtein distance to assign a score to each match.

Our pipeline first attempts to match the entity string to a concept label. If the match has a score equal to or higher than 0.7 (determined empirically), we accept that match and assign the concept ID to that entity. Otherwise, we match to a list of synonyms of ontology concepts. If that match has a score higher than the original score, we assign the ID of the matched synonym to the entity, otherwise, we revert to the original match. It is preferable to match to a concept label since these are more specific and should reflect the most common nomenclature of the concepts.

The DDI corpus has a high imbalance of positive and negative relations, which hinders the training of a classification model. Even though only entities mentioned in the same sentence are considered as candidate DDIs, there is still a ratio of 1:5.9 positive to negative instances. Other authors have suggested reducing the number of negative relations through simple rules [41, 42]. We excluded from training and automatically classify as negative the pairs that fit the following rules:

- entities have the same text (regardless of case): in nearly every case a drug does not interact with itself;
- the only text between the candidate pair is punctuation: consecutive entities, in the form of lists and enumerations, are not interacting, as well instances were the abbreviation of an entity is introduced;
- both entities have anti-positive governors: we follow the methodology proposed by [41], where the head words of entities that do not interact are used to filter less informative instances.

### BO-LSTM model

The main contribution of this work is the integration of ontology information with a neural network classification model. A domain-specific ontology is a formal definition of the concepts related to a specific subject. We can define an ontology as a tuple < *C, R* >, where C is the set of concepts and R the set of relations between the concepts, where each relation is a pair of concepts (*c*_1_, *c*_2_) with *c*_1_, *c*_2_ ∈ *E*. In our case, we consider only subsumption relations (is-a), which are transitive, i.e. if (*c*_1_, *c*_2_) ∈ *R* and (*c*_2_, *c*_3_) ∈ *R*, then we can assume that (*c*_1_, *c*_3_) is a valid relation. Then, the ancestors of concept *c* are given by

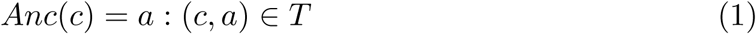

where *T* is the transitive closure of *R* on the set *E*, i.e., the smallest relation set on *E* that contains *R* and is transitive. Using this definition, we can define the common ancestors of concepts *c*_1_ and *c*_2_ as

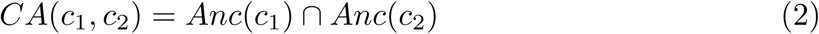

and the concatenation of the ancestors of concepts *c*_1_ and *c*_2_ as

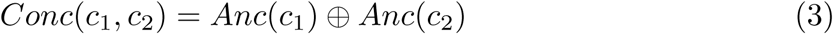

We consider two types of representations of a candidate pair based on the ancestry of its elements: the first consisting of the concatenation of the sequence of ancestors of each entity; and second, consisting of the common ancestors between both entities. Each set of ancestors is sorted by its position in the ontology so that more general concepts are in the first positions and the final position is the concept itself. Common ancestors are also used in some semantic similarity measures [43–45], since they normally represent the common information between two concepts. Due to the fact that in some cases there can be almost no overlap between the ancestors of two concepts, the concatenation provides an alternative representation.

We first represent each ontology concept as a one-hot vector *v*_*c*_, a vector of zeros except for the position corresponding to the ID of the concept. The ontology embedding layer transforms these sparse vectors into dense vectors, known as embeddings, through an embedding matrix *M* ∈ ℝ*^D×C^*, where *D* is the dimensionality of the embedding layer and *C* is the number of concepts of the ontology. Then, the output of the embedding layer is given by

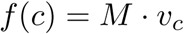

In our experiments, we set the dimensionality of the ontology embedding layer as 50, and initialized its values randomly. Then, these values were tuned during training through back-propagation.

The sequence of vectors representing the ancestors of the terms is then fed into the LSTM layer. RNNs model the probability of each element, taking into account all previous elements of the sequence. This makes sense for our objective since we want to model the semantics of each concept according to its ancestors. RNNs contain hidden units, also known as neurons, which perform linear operations on the input followed by a non-linear operation, such as sigmoid or tanh. The state of the t-th hidden unit is given by

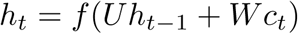

where *f* is a non-linear function, *c*_*t*_ is the element at position *t, U* is the weight matrix for the previous element and *W* the weight matrix for the current element. Since the weight matrix of each position is affected by the matrices of the previous positions, during back-propagation, the contribution of the gradient to the earlier positions may become too small or too large, leading to the vanishing and exploding gradient problems. One possible solution to these problems is through a gated RNN architecture, where the non-linear activation function is modified to regulate the contribution of earlier positions. LSTM units are a popular RNN architecture, composed of three gates (input, forget and output) and a cell, given by the following equations:

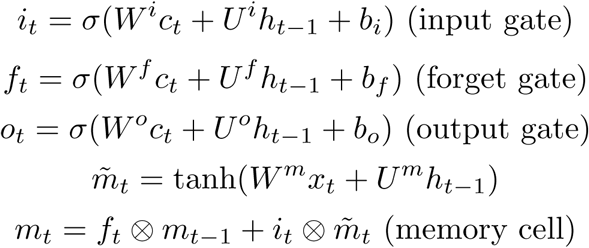

where *W*^*i*^, *W*^*f*^, *W*^*o*^ and *W^m^* correspond to the input weights of each respective gate and cell, *U*^*i*^, *U*^*f*^, *U*^*o*^, *U*^*m*^ correspond to the weights of the previous position, *σ* is the sigmoid function ⊗ and corresponds to element-wise multiplication. The output of each LSTM unit is then given by

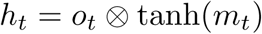

Fig 5 exemplifies how we adapted this architecture to our model, using a sequence of ontology concepts as input. After the LSTM layer, we use a max pool layer which selects the maximum *h*_*t*_ value at each position *t*. The output of the max pool layer is fed into a dense layer with a sigmoid activation function. We experimented bypassing this dense layer, obtaining inferior results. Finally, a softmax layer outputs the probability of each class.

**Fig 5.** BO-LSTM unit, using a sequence of ChEBI ontology concepts as an example. Circle refers to sigmoid function and rectangle to tanh, while “x” and “+” refer to element-wise multiplication and addition. *h*: hidden unit; 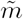: candidate memory cell; *m*: memory cell; *i* input gate; *f* forget gate; *o*: output gate;

Each configuration of our model was trained through mini-batch gradient descent with the Adam algorithm [46] and with cross-entropy as the loss function. We used the dropout strategy [47] to reduce overfitting on the trained embeddings and weights. We tuned the hyperparameters common to all configurations using only the word embeddings channel on the validation set. Each model was trained until the validation loss stopped decreasing. The experiments were performed on an Intel Xeon CPU (X3470 @ 2.93 GHz) with 16 GB of RAM.

The ChEBI and WordNet embedding layers were trained along with the other layers of the network. The DDI corpus contains 1757 of the 109k concepts of the ChEBI ontology. Since this is a relatively small vocabulary, we believe that this approach is robust enough to tune the weights. For the size of the Wordnet embedding layer, we used 50 as suggested by Xu et al. [15], while for the ChEBI embedding layer, we tested 50, 100 and 150, obtaining the best performance with 50.

### Baseline model

As a baseline, we implemented a model based on the SDP-LSTM model of Xu et al. [15] The SDP-LSTM model makes use of four types of information: Word embeddings, Part-of-speech tags, Grammatical relations and Wordnet hypernyms, which we refer to as channels. Each channel uses a specific type of input information to train an LSTM-based RNN layer, which is then connected to a max pooling layer, the output of the channel. The output of each channel is concatenated, and connected to a densely-connected hidden layer, with a sigmoid activation function, while a softmax layer outputs the probabilities of each class.

Xu et. al show that it is possible to obtain high performance on a relation extraction task using only the word representations channel. For this reason, we use a version of our model with only this channel as the baseline. We employ the previously mentioned pre-trained word embeddings as input to the LSTM layer.

Additionally, we make use of Wordnet as an external source of information. The authors of the SDP-LSTM model showed that Wordnet contributed to an improvement of the F1-score on a relation extraction task. We use the tool developed by Ciaramita and altun [39] to obtain the Wordnet classes of each word according to 41 semantic categories, such as “noun.group” and “verb.change”. The embeddings of this channel were set to be 50-dimensional and tuned during the training of the model.

## References

1. Zelenko D, Zelenko D, Aone C, Aone C, Richardella A, Richardella A. Kernel Methods for Relation Extraction. Journal of Machine Learning Research. 2003;3:1083–1106. doi:10.3115/1118693.1118703.

2. Reichartz F, Korte H, Paass G. Semantic relation extraction with kernels over typed dependency trees. Proceedings of the 16th ACM SIGKDD international conference on Knowledge discovery and data mining - KDD ‘10. 2010; p. 773. doi:10.1145/1835804.1835902.

3. Kambhatla N. Combining lexical, syntactic, and semantic features with maximum entropy models for extracting relations. Proceedings of the ACL 2004 on Interactive poster and demonstration sessions. 2004; p. 22. doi:10.3115/1219044.1219066.

4. Collobert R, Weston J, Bottou L, Karlen M, Kavukcuoglu K, Kuksa P. Natural language processing (almost) from scratch. Journal of Machine Learning Research. 2011;12(Aug):2493–2537.

5. Erhan D, Bengio Y, Courville A, Manzagol PA, Vincent P, Bengio S. Why Does Unsupervised Pre-training Help Deep Learning? J Mach Learn Res. 2010;11:625–660.

6. Mikolov T, Sutskever I, Chen K, Corrado GS, Dean J. Distributed Representations of Words and Phrases and their Compositionality. In: Burges CJC, Bottou L, Welling M, Ghahramani Z, Weinberger KQ, editors. Advances in Neural Information Processing Systems 26. Curran Associates, Inc.; 2013. p. 3111–3119. Available from: http://papers.nips.cc/paper/5021-distributed-representations-of-words-and-phrases-and-their-comppdf.

7. Pennington J, Socher R, Manning CD. GloVe: Global Vectors for Word Representation. In: Empirical Methods in Natural Language Processing (EMNLP); 2014. p. 1532–1543. Available from: http://www.aclweb.org/anthology/D14-1162.

8. Huang CC, Lu Z. Community challenges in biomedical text mining over 10 years: Success, failure and the future. Briefings in Bioinformatics. 2016;17(1):132–144. doi:10.1093/bib/bbv024.

9. Al Hamid A, Ghaleb M, Aljadhey H, Aslanpour Z. A systematic review of hospitalization resulting from medicine-related problems in adult patients. British Journal of Clinical Pharmacology. 2014;78(2):202–217. doi:10.1111/bcp.12293.

10. Bouvy JC, De Bruin ML, Koopmanschap MA. Epidemiology of adverse drug reactions in Europe: a review of recent observational studies. Drug safety. 2015;38(5):437–453.

11. Dou D, Wang H, Liu H. Semantic data mining: A survey of ontology-based approaches. In: Proceedings of the 2015 IEEE 9th International Conference on Semantic Computing (IEEE ICSC 2015); 2015. p. 244–251.

12. Hochreiter S, Schmidhuber J. Long short-term memory. Neural computation. 1997;9(8):1735–1780.

13. Miwa M, Bansal M. End-to-end Relation Extraction using LSTMs on Sequences and Tree Structures. Acl 2016. 2016; p. 10. doi:10.18653/v1/P16-1105.

14. Zhang S, Zheng D, Hu X, Yang M. Bidirectional long short-term memory networks for relation classification. In: Proceedings of the 29th Pacific Asia Conference on Language, Information and Computation; 2015. p. 73–78.

15. Xu Y, Mou L, Li G, Chen Y. Classifying Relations via Long Short Term Memory Networks along Shortest Dependency Paths. In: In Proceedings of Conference on Empirical Methods in Natural Language Processing. September; 2015. p. 1785–1794.

16. Smith B, Ashburner M, Rosse C, Bard J, Bug W, Ceusters W, et al. The OBO Foundry: coordinated evolution of ontologies to support biomedical data integration. Nature biotechnology. 2007;25(11):1251.

17. Hastings J, De Matos P, Dekker A, Ennis M, Harsha B, Kale N, et al. The ChEBI reference database and ontology for biologically relevant chemistry: Enhancements for 2013. Nucleic Acids Research. 2013;41(D1):456–463. doi:10.1093/nar/gks1146.

18. Ferreira JD, In´acio B, Salek RM, Couto FM. Assessing Public Metabolomics Metadata, Towards Improving Quality. Journal of integrative bioinformatics. 2017;14(4).

19. Li Q, Li T, Chang B. Learning Word Sense Embeddings from Word Sense Definitions. In: Lin CY, Xue N, Zhao D, Huang X, Feng Y, editors. Natural Language Understanding and Intelligent Applications. Cham: Springer International Publishing; 2016. p. 224–235.

20. Ma N, B HtZ, Xiao X. An Ontology-Based Latent Semantic Indexing Approach Using Long Short-Term Memory Networks. Web and Big Data. 2017;10366(2):185–199. doi:10.1007/978-3-319-63579-8.

21. Goyal P, Ferrara E. Graph embedding techniques, applications, and performance: A survey. arXiv preprint arXiv:170502801. 2017;.

22. Kong X, Cao B, Yu PS. Multi-label Classification by Mining Label and Instance Correlations from Heterogeneous Information Networks. In: Proceedings of the 19th ACM SIGKDD International Conference on Knowledge Discovery and Data Mining. KDD ‘13. New York, NY, USA: ACM; 2013. p. 614–622. Available from: http://doi.acm.org/10.1145/2487575.2487577.

23. Dasigi P, Ammar W, Dyer C, Hovy E. Ontology-Aware Token Embeddings for Prepositional Phrase Attachment. In: Proceedings of the 55th Annual Meeting of the Association for Computational Linguistics (Volume 1: Long Papers). Association for Computational Linguistics; 2017. p. 2089–2098. Available from: http://www.aclweb.org/anthology/P17-1191.

24. Mu¨ller HMM, Kenny EE, Sternberg PW. Textpresso: an ontology-based information retrieval and extraction system for biological literature. PloS Biology. 2004;2(11):e309. doi:10.1371/journal.pbio.0020309.

25. Lamurias A, Ferreira JD, Couto FM. Identifying interactions between chemical entities in biomedical text. Journal of integrative bioinformatics. 2014;11(3):1–16.

26. Tripodi I, Boguslav M, Haylu N, Hunter LE. Knowledge-base-enriched relation extraction. In: Proceedings of the Sixth BioCreative Challenge Evaluation Workshop. Bethesda, MD USA. vol. 1; 2017. p. 163–166.

27. Herrero-Zazo M, Segura-Bedmar I, Mart´ınez P, Declerck T. The DDI corpus: An annotated corpus with pharmacological substances and drug-drug interactions. Journal of Biomedical Informatics. 2013;46(5):914–920. doi:10.1016/j.jbi.2013.07.011.

28. Wheeler DL, Barrett T, Benson DA, Bryant SH, Canese K, Chetvernin V, et al. Database resources of the national center for biotechnology information. Nucleic acids research. 2006;35(suppl_1):D5–D12.

29. Segura-Bedmar I, Mart´ınez P, Herrero-Zazo M. Lessons learnt from the DDIExtraction-2013 Shared Task. Journal of Biomedical Informatics. 2014;51(May):152–164. doi:10.1016/j.jbi.2014.05.007.

30. Zhao Z, Yang Z, Luo L, Lin H, Wang J. Drug drug interaction extraction from biomedical literature using syntax convolutional neural network. Bioinformatics. 2016;32(November):btw486. doi:10.1093/bioinformatics/btw486.

31. Sahu SK, Anand A. Drug-Drug Interaction Extraction from Biomedical Text Using Long Short Term Memory Network. CEUR Workshop Proceedings. 2017;1828:53–59. doi:10.1145/2910896.2910898.

32. Ashburner M, Ball CA, Blake JA, Botstein D, Butler H, Cherry JM, et al. Gene Ontology: tool for the unification of biology. Nature genetics. 2000;25(1):25.

33. Dritsou V, Topalis P, Mitraka E, Dialynas E, Louis C. miRNAO: An Ontology Unfolding the Domain of microRNAs. In: IWBBIO; 2014. p. 989–1000.

34. Langlotz CP. RadLex: a new method for indexing online educational materials; 2006.

35. Rath A, Olry A, Dhombres F, Brandt MM, Urbero B, Ayme S. Representation of rare diseases in health information systems: the Orphanet approach to serve a wide range of end users. Human mutation. 2012;33(5):803–808.

36. Kibbe WA, Arze C, Felix V, Mitraka E, Bolton E, Fu G, et al. Disease Ontology 2015 update: an expanded and updated database of human diseases for linking biomedical knowledge through disease data. Nucleic acids research. 2014;43(D1):D1071–D1078.

37. Ou M, Cui P, Wang F, Wang J, Zhu W. Non-transitive hashing with latent similarity components. In: Proceedings of the 21th ACM SIGKDD International Conference on Knowledge Discovery and Data Mining. ACM; 2015. p. 895–904.

38. Lamurias A, Ferreira J, Couto F. Improving chemical entity recognition through h-index based semantic similarity. Journal of Cheminformatics. 2015;7(Suppl 1):S13, 1–20. DOI:https://doi.org/10.1186/1758-2946-7-S1-S13.

39. Ciaramita M, Altun Y. Broad-coverage sense disambiguation and information extraction with a supersense sequence tagger. In: Proceedings of the 2006 Conference on Empirical Methods in Natural Language Processing. Association for Computational Linguistics; 2006. p. 594–602.

40. Pyysalo S, Ginter F, Moen H, Salakoski T, Ananiadou S. Distributional Semantics Resources for Biomedical Text Processing. Proceedings of LBM 2013. 2013;.

41. Chowdhury MFM, Lavelli A. FBK-irst: A multi-phase kernel based approach for drug-drug interaction detection and classification that exploits linguistic information. Atlanta, Georgia, USA. 2013;351:53.

42. Kim S, Liu H, Yeganova L, Wilbur WJ. Extracting drug–drug interactions from literature using a rich feature-based linear kernel approach. Journal of biomedical informatics. 2015;55:23–30.

43. Resnik P. Using information content to evaluate semantic similarity in a taxonomy. In: International Joint Conference on Artificial Intelligence. vol. 14. Citeseer; 1995. p. 448–453.

44. Jiang JJ, Conrath DW. Semantic Similarity Based on Corpus Statistics and Lexical Taxonomy. CoRR. 1997;cmp-lg/9709008.

45. Lin D. An Information-Theoretic Definition of Similarity. In: Proceedings of the Fifteenth International Conference on Machine Learning. ICML ‘98. San Francisco, CA, USA: Morgan Kaufmann Publishers Inc.; 1998. p. 296–304. Available from: http://dl.acm.org/citation.cfm?id=645527.657297.

46. Kingma DP, Ba J. Adam: A Method for Stochastic Optimization. CoRR. 2014;abs/1412.6980.

47. Hinton GE, Srivastava N, Krizhevsky A, Sutskever I, Salakhutdinov R. Improving neural networks by preventing co-adaptation of feature detectors. CoRR. 2012;abs/1207.0580.

